# Widely targeted metabolomics analysis reveals differences in volatile metabolites among four *Angelica* species

**DOI:** 10.1101/2022.12.01.518649

**Authors:** Jiaojiao Ji, Lanlan Zang, Tingting Lu, Cheng Li, Xiaoxu Han, Soo-Rang Lee, Li Wang

## Abstract

*Angelica* L. has attracted global interest for its traditional medicinal uses and commercial values. However, few studies have focused on the metabolomic differences among the *Angelica* species. In this study, employing the widely targeted metabolomics based on gas chromatography-tandem mass spectrometry, the metabolomes of four *Angelica* species were analyzed (*Angelica sinensis* (Oliv.) Diels (*A. sinensis*), *Angelica biserrata* (R.H.Shan & Yuan) C.Q.Yuan & R.H.Shan (*A. biserrata*), *Angelica dahurica* (Hoffm.) Benth. & Hook.f. ex Franch. & Sav. (*A. dahurica*), *Angelica keiskei* Koidz. (*A. keiskei*)). A total of 698 volatile metabolites were identified and classified into fifteen different categories. The metabolomic analysis indicated that 7-hydroxycoumarin and Z-ligustilide were accumulated at significantly higher levels in *A. sinensis*, whereas the opposite pattern was observed for bornyl acetate. In addition, a high correspondence between the dendrogram of metabolite contents and phylogenetic positions was detected in the four species. This study provides a biochemical map for the exploitation, application and development of the *Angelica* species as medicinal plants or health-related dietary supplements.

## 1. Introduction

*Angelica* L., a genus in the family Apiaceae, is comprised of 90 species of herbs that are widespread in north-temperate regions, especially Eurasia [1,2]. Many plants in the genus have long been used in traditional Chinese medicine (TCM) [3], in particular, the dried roots of *Angelica* have been widely used for nourishing blood, regulating menstruation, and analgesic [1,4]. Various herbal preparations containing *Angelica* species are available over the counter, not only in China, but also in Europe and American countries [5,6]. Besides its medicinal value, *Angelica* is also highly appreciated in various industrial applications such as the dietary supplements, perfumery, and cosmetics [1,7,8]. A previous study demonstrated that the pharmacological activity of aromatic and medicinal plants is attributed to its effective volatile components [9]. Plants in *Angelica* are extremely rich in secondary metabolites, including coumarins, flavonoids, terpenoids, as well as volatiles oils (VOs) [1,3]. Modern medical research has revealed that the VOs composition is mainly responsible for the medicinal properties of the genus *Angelica* [10]. VOs are complex mixture of low molecular weight volatile compounds that are isolated from the raw plant material by distillation [11], which have been reported to treat serious health diseases, involving gynecological diseases, fever, and arthritis [1,12]. There are a couple of good examples showing the proven effects of VOs in *Angelica* species. Phthalides of *Angelica sinensis* (Oliv.) Diels (*A. sinensis*) are one of the highly effective VOs to analgesic and sedative activities [6,13]. *Angelica biserrata* (R.H.Shan & Yuan) C.Q.Yuan & R.H.Shan (*A. biserrata*) also contains active ingredients such as oxygenates, terpenoids, ketones and esters with analgesic and anti-inflammatory effects [14]. However, most of current studies only focused on several targeted compounds in a single *Angelica* species. There have been no comprehensive and comparative studies examining the volatile metabolites of multiple *Angelica* species. It has posed a major obstacle to the application and exploitation of the medicinal plants in *Angelica* species.

With the development of metabolomics, high-throughput and high-resolution methods such as headspace solid phase micro-extraction gas chromatography-mass spectrometry (HS-SPME-GC-MS) have been widely used to identify metabolite profiles and detect differences in the biochemical compositions of aromatic and medicinal plants [10,15,16]. The four species *A. biserrata*, *Angelica dahurica* (Hoffm.) Benth. & Hook.f. ex Franch. & Sav. (*A. dahurica*), *Angelica keiskei* Koidz. (*A. keiskei*) and *A. sinensis* are the representative medicinal plants in *Angelica*, and it is noteworthy that roots of *A. sinensis* are one of the most widely prescribed medicine in China owing to its rich VOs [6]. In this study, volatile metabolites of four *Angelica* species were identified and quantified using widely targeted metabolomics. The aim was to reveal the differed accumulation of medicinally important metabolites among the four species. This study provides useful information for the chemical composition of *Angelica* plants and may help the identification of the biologically active substances responsible for the pharmacological activity of *Angelica* plants.

## 2. Materials and methods

### 2.1. Plant samples

Four species in genus *Angelica*, including *A. sinensis*, *A. dahurica*, *A. biserrata*, and *A. keiskei*, were analyzed in this study. The *A. sinensis* plants were collected from Minxian County, Gansu Province, China. The *A. dahurica*, *A. biserrata*, and *A. keiskei* plants were collected from Shenzhen City, Guangdong Province, China. The specimens of the four species were deposited at Agricultural Genomics Institute at Shenzhen, Chinese Academy of Agricultural Sciences and identified by Prof. Li Wang (*A. sinensis* (202011002), *A. dahurica* (202008001), *A. biserrata* (202106003) and *A. keiskei* (202109004)).

Roots of each species were sampled with three biological replicates. The collected roots were washed, naturally dried, frozen in liquid nitrogen, and then stored at -80□ for further analysis.

### 2.2. Solid phase microextraction (SPEM) extraction

The samples were ground into powder in liquid nitrogen. Powdered samples (1 g) were weighed and transferred immediately to a 20 mL head-space vial (Agilent, Palo Alto, CA, USA), containing NaCl saturated solution to inhibit potential enzyme reactions. The headspace vials were sealed using crimp-top caps. As for SPME analysis, each vial was placed in 60□ for 5 min, and then a 120 µm DVB/CWR/PDMS fiber (Agilent, Palo Alto, CA, USA) was exposed to the headspace of the sample for 15 min at 100□. DVB/CWR/PDMS are three-phase fiber heads, which were confirmed to be able to extract more volatile metabolites than other fiber headers [17]. The quality control (mix) sample was prepared by mixing equal volumes of samples into a single tube, and during the instrumental analysis, a quality control sample was inserted into each of the 10 test samples to check the repeatability of the analysis process (Figure S1 and Table S1).

### 2.3. GC-MS analysis

After the extraction procedure, the fiber was transferred to the injection port of the GC-MS system (Model 8890; Agilent, Palo Alto, CA, USA). The SPME fiber was desorbed and maintained in the injection port at 250□ for 5 min in the split-less mode. The identification and quantification of volatile metabolites was carried out using an Agilent Model 8890 GC and a 7000 D mass spectrometer (Agilent, Palo Alto, CA, USA), equipped with a 30 m × 0.25 mm × 0.25 μm DB-5MS (5% phenyl-polymethylsiloxane) capillary column. Helium was used as the carrier gas at a linear velocity of 1.2 mL/min. The injector temperature was kept at 250□ and the detector at 280□. The oven temperature was programmed as followings: 40□ (3.5 min), increasing at 10□ min^-1^ to 100□, 7□ min^-1^ to 180□, 25□ min^-1^ to 280□ and hold for 5 min. Mass spectra was recorded in electron impact ionization mode at 70 eV. The quadrupole mass detector, ion source and transfer line temperatures were set, respectively, at 150, 230 and 280□. For the identification and quantification of analytes, the MS was selected ion monitoring mode.

### 2.4. Qualitative and quantitative analysis

After the mass spectrometry analysis, all raw data were analyzed with the software Qualitative Analysis Workflows B.08.00 (Agilent, Palo Alto, CA, USA). The qualitative analysis of primary and secondary mass spectrometry data was annotated based on the self-built database MWDB (Metware Biotechnology Co., Ltd. Wuhan, China) and the publicly available metabolite databases.

### 2.5 Statistical analysis

After the metabolite data was transformed with Hellinger transformation, principal component analysis (PCA) was performed using the function rda in the R package vegan v 2.6-2 [18]. In addition, the data set that was log_10_ transform and mean centering were imported into the R package MetaboAnalystR v5.0 [19] to conduct orthogonal partial least squares-discriminant analysis (OPLS-DA) and extract the variable important in projection (VIP) value from the analysis results to evaluate the variance importance of compounds. The values of R2X, R2Y, and Q2 for OPLS-DA models were showed in Figure S2. In order to avoid overfitting, a permutation test (100 permutations) was performed. Based on Bray-Curtis’s dissimilarity distances of the composition and abundance of all volatile metabolites after ln transform, which were calculated using the function vegdist built in vegan, hierarchical clustering was visualized with the R package factoextra v.1.0.7 [20].

The chloroplast sequence alignments of *A. sinensis*, *A. dahurica*, *A. biserrata*, and *A. keiskei* were generated using MAFFT v7.475 [21]. Phylogenetic trees were constructed by maximum likelihood using IQ-TREE v 2.1.2 [22] with *Hydrocotyle sibthorpioides* Lam. as an outgroup.

All identified metabolites were annotated with KEGG database (http://www.kegg.jp/kegg/compound/) and further subjected to KEGG enrichment analyses with the R package clusterProfiler v. 4.4.4 [23].

## 3. Results

### 3.1. Metabolomics profiling of four Angelica species

Widely targeted metabolomics offered a promising way for the chemical screening of volatile metabolites and allowed the characterization of new metabolites in *Angelica* [10]. In this study, to get insight into differences of volatile metabolites among four *Angelica* species, the root metabolomics data were generated. A total of 698 non-redundant volatile metabolites were qualified and quantified based on GC-MS (Table S1). Among them, 616, 536, 576, and 545 volatile metabolites in *A. sinensis*, *A. dahurica*, *A. biserrata*, *A. keiskei*, respectively. Three hundred and ninety-one metabolites were commonly detected in the roots of all four species (Figure 1a).

**Figure 1.**
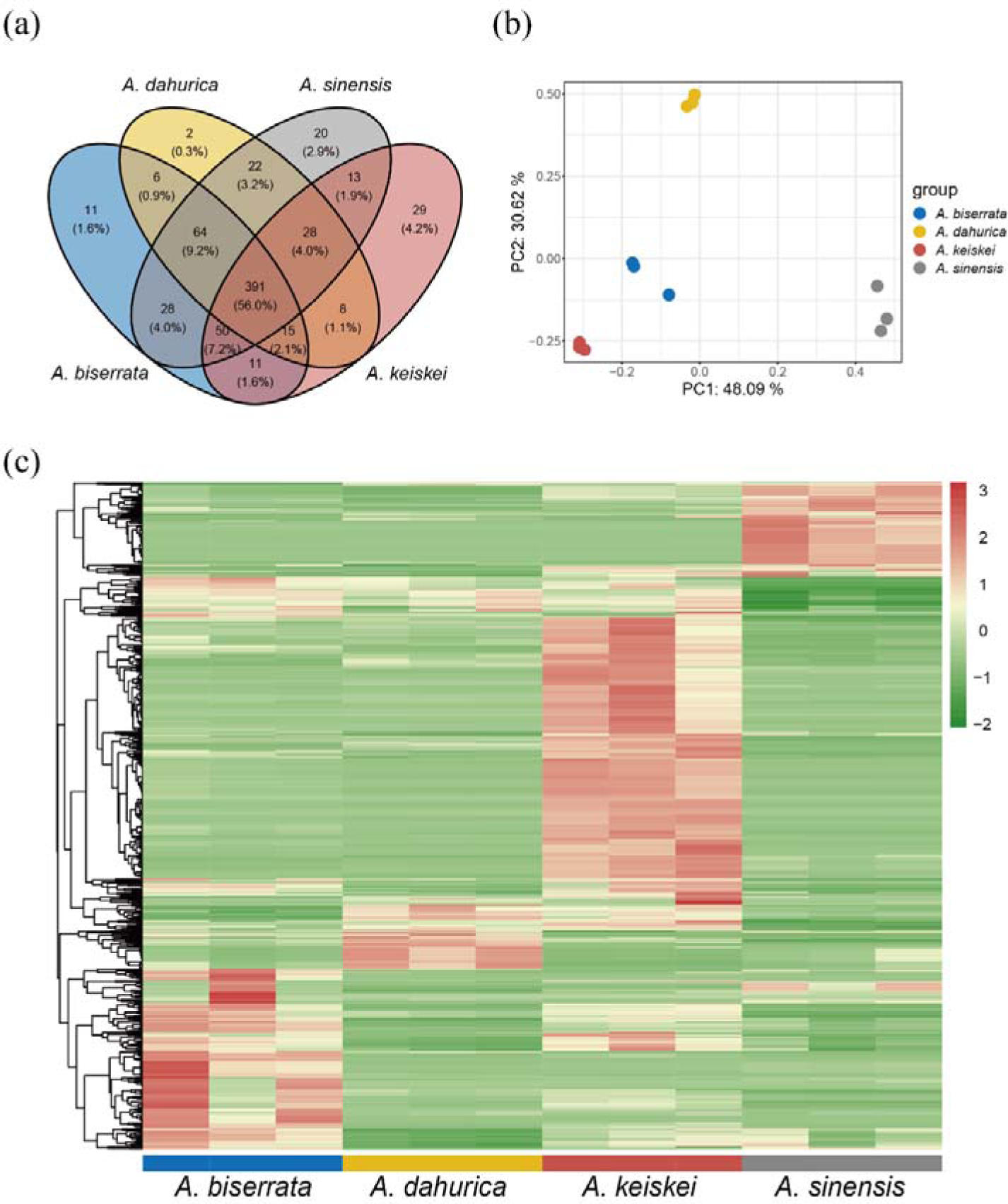
An overview of volatile metabolites among four *Angelica* species. (a) Venn diagram showing the number of common and specific metabolites in the four species. (b) PCA of volatile metabolites for the four species with three biological replicates. (c) Heatmap clustering of volatile metabolites identified from the four species. Volatile metabolite abundance was *Z-*score transformed. The color-coded scale grading from green to red corresponds to the content of volatile metabolites shifting from low to high.

PCA of the metabolome data, transformed with Hellinger transformation method, the samples were divided into four distinct groups corresponding to the four species. And the three biological replicates were clustered together, suggesting that the data are reproducible and reliable. Based on the PCA plot, where PC1 and PC2 explained 48.09% and 30.62% of the total variance, respectively. Of the four clusters, PC1 mainly differentiated *A. sinensis* from the other *Angelica* species, while PC2 primarily segregated *A. dahurica* from the other *Angelica* species (Figure 1b).

The abundances of volatile metabolites were transformed by *Z*-score and then subjected to hierarchical clustering analysis (Figure 1c). The results showed that showed significant differences among the four species. Of these, the abundance of volatile metabolites in *A. keiskei* was the highest.

### 3.2. Identification of differential metabolites in four Angelica species

To explore the metabolite composition of the four species, 698 volatile metabolites were classified into 15 different categories, including terpenoids, ester, heterocyclic, aromatics and 11 others (Figure 2). The terpenoids took up the highest proportion of all measured volatile metabolites in the four *Angelica* species, followed by heterocyclic compounds, eater, and aromatics. Notably, *A. sinensis* contained a relatively lower proportion (43.34%) of terpenoids than other *Angelica* species, but exhibited a more balanced metabolite composition in volatile metabolites. In contrast, the amount of terpenoids accounted for more than half of the total volatile metabolites in *A. dahurica*, *A. biserrata*, and *A. keiskei*, especially in *A. dahurica*, its proportion reached up to 75.64%.

**Figure 2.**
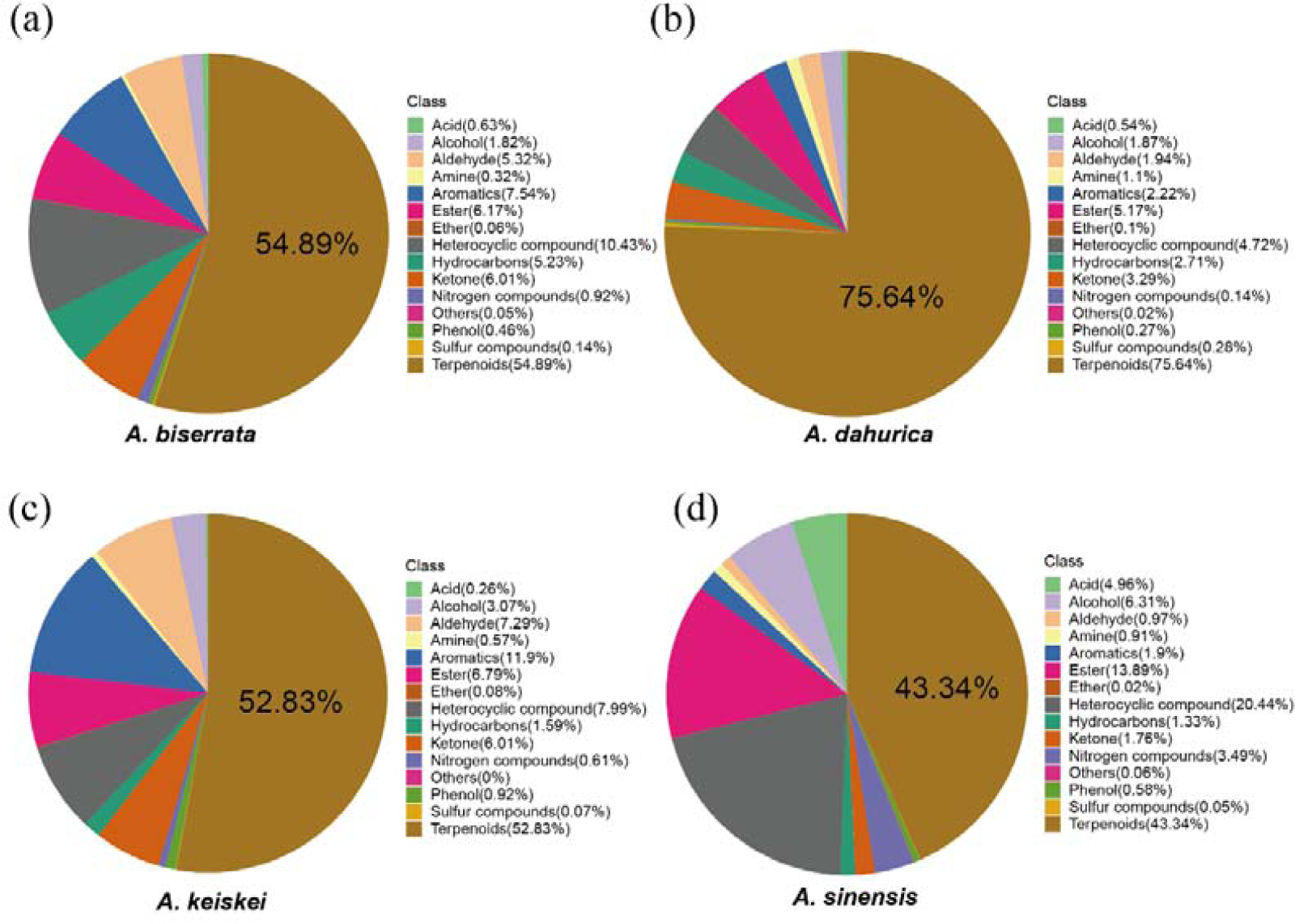
Classification and proportion of volatile metabolites detected in the four *Angelica* species. (a) *A. biserrata*, (b) *A. dahurica*, (c) *A. keiskei*, (d) *A. sinensis*.

In addition, the relative abundance of each metabolite category in the four species were compared with Kruskal-Wallis test. The *p* values corrected by the bonferroni method showed significant difference in eight categories among the four species, including alcohol, aldehyde, aromatics, ester, heterocyclic compounds, hydrocarbons, ketone and terpenoids (Figure 3, Figure S3; *p*-values were shown in the Table S2). Terpenoids was the most different metabolites among the four species. In each of the eight categories, *A. sinensis* and *A. keiskei* were significantly different from at least one species. These results indicated that the difference between the presented *Angelica* species lies in the representation of individual components in the metabolomic profiles of the samples, while the qualitative composition is approximately the same.

**Figure 3.**
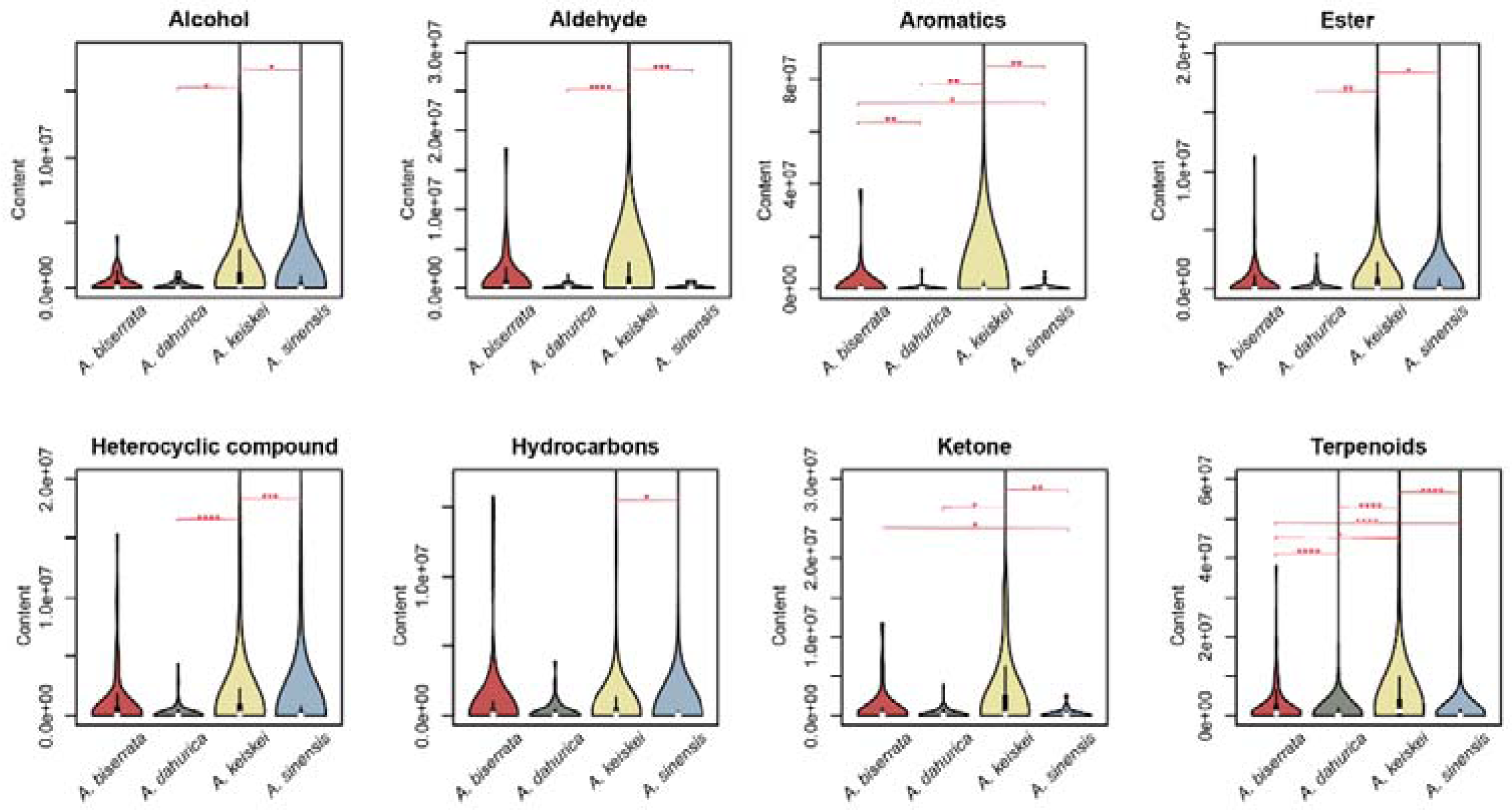
Comparison for the relative abundance of seven categories (alcohol, aromatics, aldehyde, ester, heterocyclic compounds, ketone and terpenoids) with significant differences in the four *Angelica* species.

Here, this study performed hierarchical clustering analysis based on Bray-Curtis’s dissimilarity distances of the composition and abundance of volatile metabolites in the four *Angelica* species. The dendrogram (Figure 4a) showed high correspondence with the phylogenetic tree (Figure 4b) based on chloroplast sequences, suggesting a correlation relationship between the volatile metabolites and the phylogenetic relationships.

**Figure 4.**
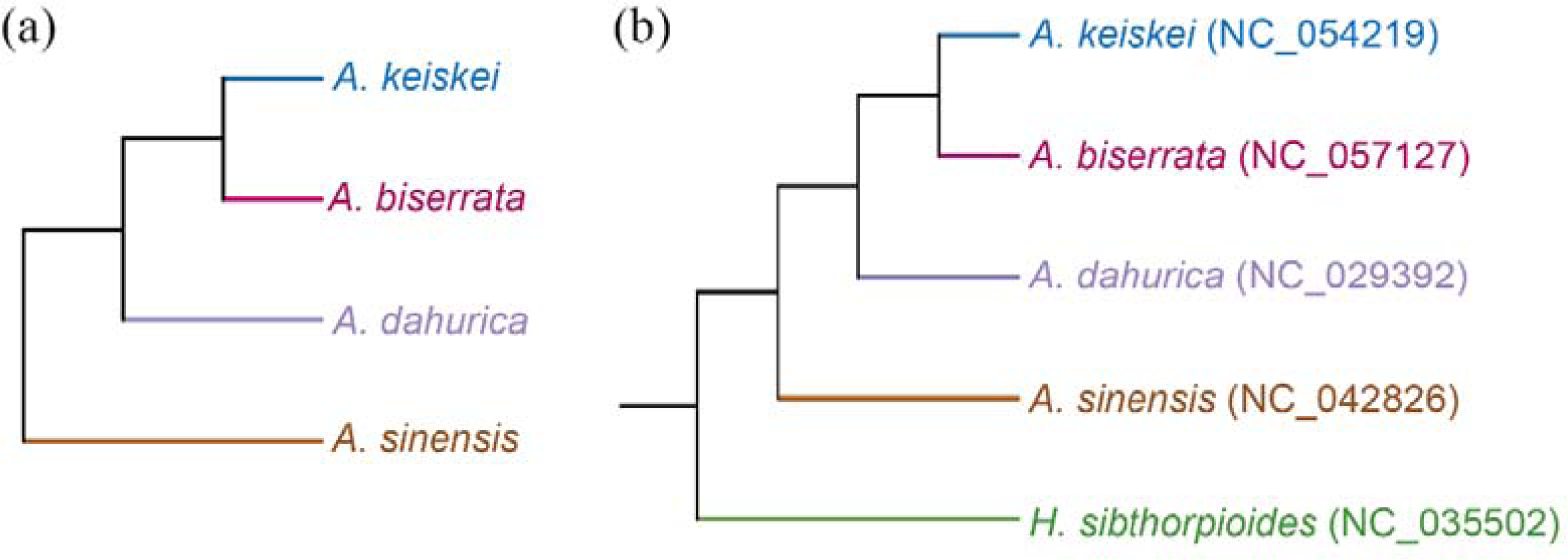
Hierarchical clustering based on the similarity of volatile metabolites (a) and phylogenetic tree of the four *Angelica* species and *H. sibthorpioides* (b). The chloroplast sequences above were available in GenBank of NCBI.

### 3.3. Differential metabolites between A. sinensis and the three other Angelica species

To further identify the metabolites responsible for differences among the four *Angelica* species, significantly different accumulated metabolites between groups were screened by |Log2FC|≥1 and VIP≥1. *A. sinensis* belongs to *Sinodielsia* clade in *Angelica* genus, that was phylogenetically distant from core *Angelica* group, including *A. biserrata, A. dahurica*, *A. keiskei* [2,24]. Moreover, PC1 mainly differentiated *A. sinensis* from the other three species (Figure 1b), therefore, *A. sinensis* was used to comparison to other three species. Interestingly, there were fewer up-regulated metabolites in *A. sinensis* when compared with the other species. And no significantly enriched pathway was detected in the KEGG enrichment results of these differential metabolites, which could be a bias caused by the small dataset. Compared with *A. biserrata*, 446 significantly differential metabolites (123 up-regulated and 323 down-regulated) were screened in *A. sinensis* (Figure 5a), and the top 3 enrichment pathways of these substance were metabolic pathways (23 metabolites with *p* = 0.19), tyrosine metabolism (3 metabolites with *p* = 0.22) and limonene and pinene degradation (5 metabolites with *p* = 0.24) (Figure 5d). Compared with *A. dahurica*, 429 significantly differential metabolites (169 up-regulated and 260 down-regulated) were detected in *A. sinensis* (Figure 5b), and the top 3 enrichment pathways of these metabolites were tyrosine metabolism (3 metabolites with *p* = 0.21), limonene and pinene degradation (5 metabolites with *p* = 0.22) and metabolic pathways (22 metabolites with *p* = 0.26) (Figure 5e). When compared with *A. keiskei*, 502 significantly differential metabolites (105 up-regulated and 397 down-regulated) were identified in *A. sinensis* (Figure 5c), which were the most abundant compared with the other two group, and the top 3 enrichment pathways of these metabolites were metabolic pathways (25 metabolites with *p* = 0.09), biosynthesis of various plant secondary metabolites (5 metabolites with *p* = 0.10), and tyrosine metabolism (3 metabolites with *p* = 0.26) (Figure 5f).

**Figure 5.**
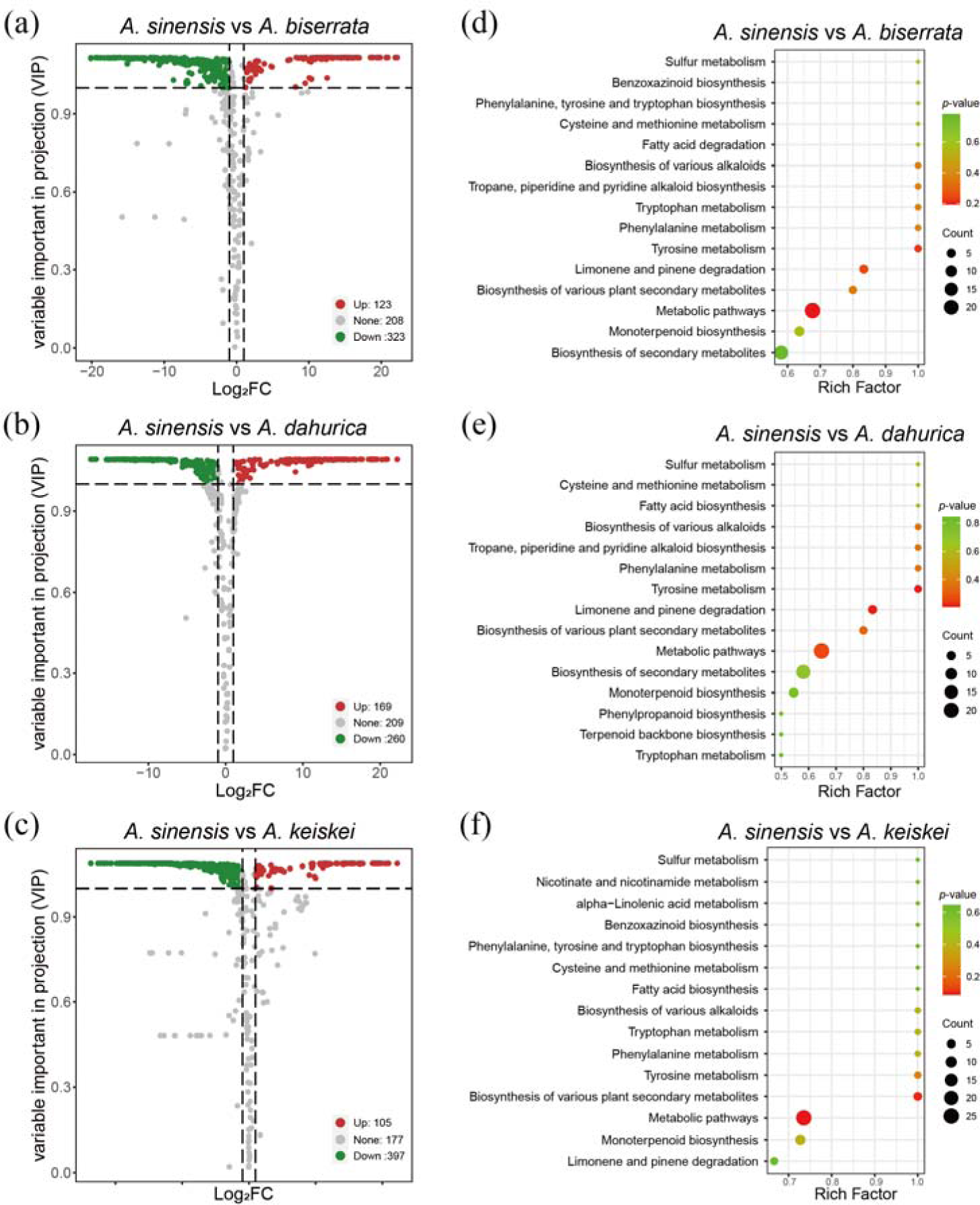
The overall distribution and KEGG enrichment analysis of differential metabolites between *A. sinensis* and the three other *Angelica* species. (a-c) Volcano plots for differential metabolites between *A. sinensis* and the three other *Angelica* species. (a) *A. sinensis* vs *A. biserrata*. (b) *A. sinensis* vs *A. dahurica*. (c) *A. sinensis* vs *A. keiskei*. Colors of metabolites indicated significant differences (red, upregulated; green, downregulated). (d-f) KEGG pathway enrichment analysis of differential metabolites for *A. sinensis* vs *A. biserrata* (d), *A. sinensis* vs *A. dahurica* (e) and *A. sinensis* vs *A. keiskei* (f). Color of the bubbles represented statistical significance of the enriched terms, and the size of the bubbles represented number of differentially enriched metabolites. The pathway of “Biosynthesis of various plant secondary metabolites” including: crocin biosynthesis, cannabidiol biosynthesis, mugineic acid biosynthesis, pentagalloylglucose biosynthesis, benzoxazinoid biosynthesis, gramine biosynthesis, coumarin biosynthesis, furanocoumarin biosynthesis, hordatine biosynthesis, podophyllotoxin biosynthesis.

In order to delve into the details of the volatile metabolite difference between *A. sinensis* and the other three species, the most significantly twenty metabolites (the top 10 for up-regulation and down-regulation, respectively) were selected (Figure 6). It was discovered that hippuric acid, 7-hydroxycoumarin and 7-ethoxycoumarin were more enriched in *A. sinensis* than the three other *Angelica* species. In addition, the abundance of 3-butylisobenzofuran-1(3H)-one in *A. sinensis* was also substantially higher than that in *A. dahurica and A. keiskei* (log_2_FC > 19). Meanwhile, the metabolites γ-terpinene and bornyl acetate in *A. dahurica, A. keiskei* and *A. biserrata* were in high abundance, but the metabolites were lower in *A. sinensis*.

**Figure 6.**
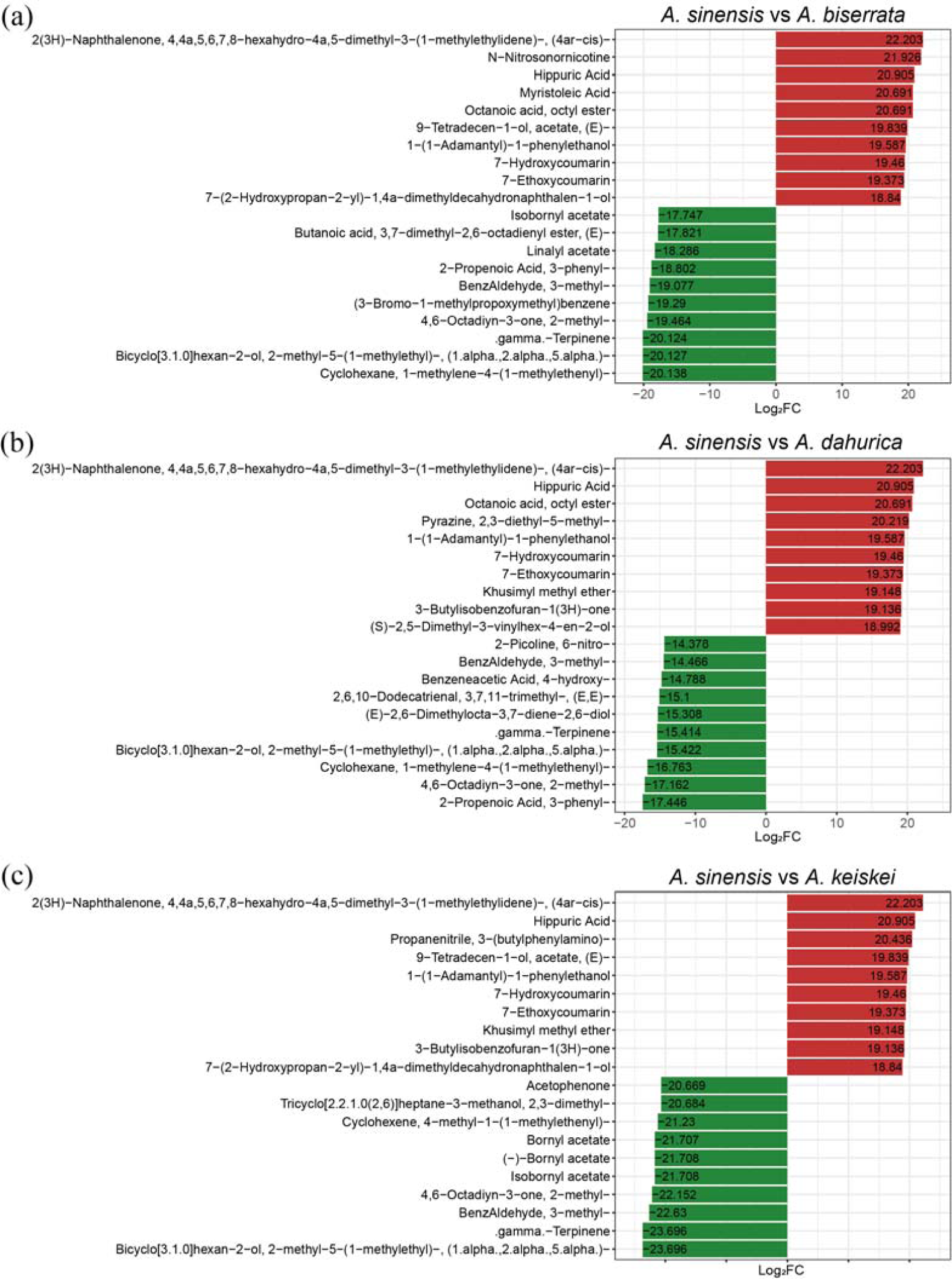
The top 20 metabolites of significantly differential volatiles between *A. sinensis* and three other *Angelica* species. Red indicates the more abundant metabolites in *A. sinensis* compared to *A. biserrata* (a), *A. dahurica* (b), *A. keiskei* (c). Green indicates the lower levels of metabolites in *A. sinensis* than that in other species.

### 3.4. Differential metabolites between A. keiskei and the three other Angelica species

Given the abundance of volatile metabolites in the root of the *A. keiskei* was the highest among the four species. It showed that the non-medicinal parts of *A. keiskei* also have potential to be exploited for practical uses. Therefore, the differences of metabolites between *A. keiskei* and the other *Angelica* species were further compared. The volcanic map visually showed the overall distribution of differential metabolites in each comparison. Four hundred and one significantly different metabolites (308 up-regulated and 93 down-regulated) were detected in the comparison between *A. keiskei* and *A. biserrata* (Figure 7a), which were related to phenylpropanoid biosynthesis (2 metabolites with *p* = 0.21), metabolic pathways (17 metabolites with *p* = 0.38) and tyrosine metabolism (3 metabolites with *p* = 0.45) (Figure 7c). Four hundred seventy-three significantly different metabolites (421 up-regulated and 52 down-regulated) were detected in the comparison between *A. keiskei* and *A. dahurica* (Figure 7b), which were associated with sesquiterpenoid and triterpenoid biosynthesis (8 metabolites with *p* = 0.14), monoterpenoid biosynthesis (9 metabolites with *p* = 0.23) and biosynthesis of secondary metabolites (22 metabolites with *p* = 0.39) (Figure 7d).

**Figure 7.**
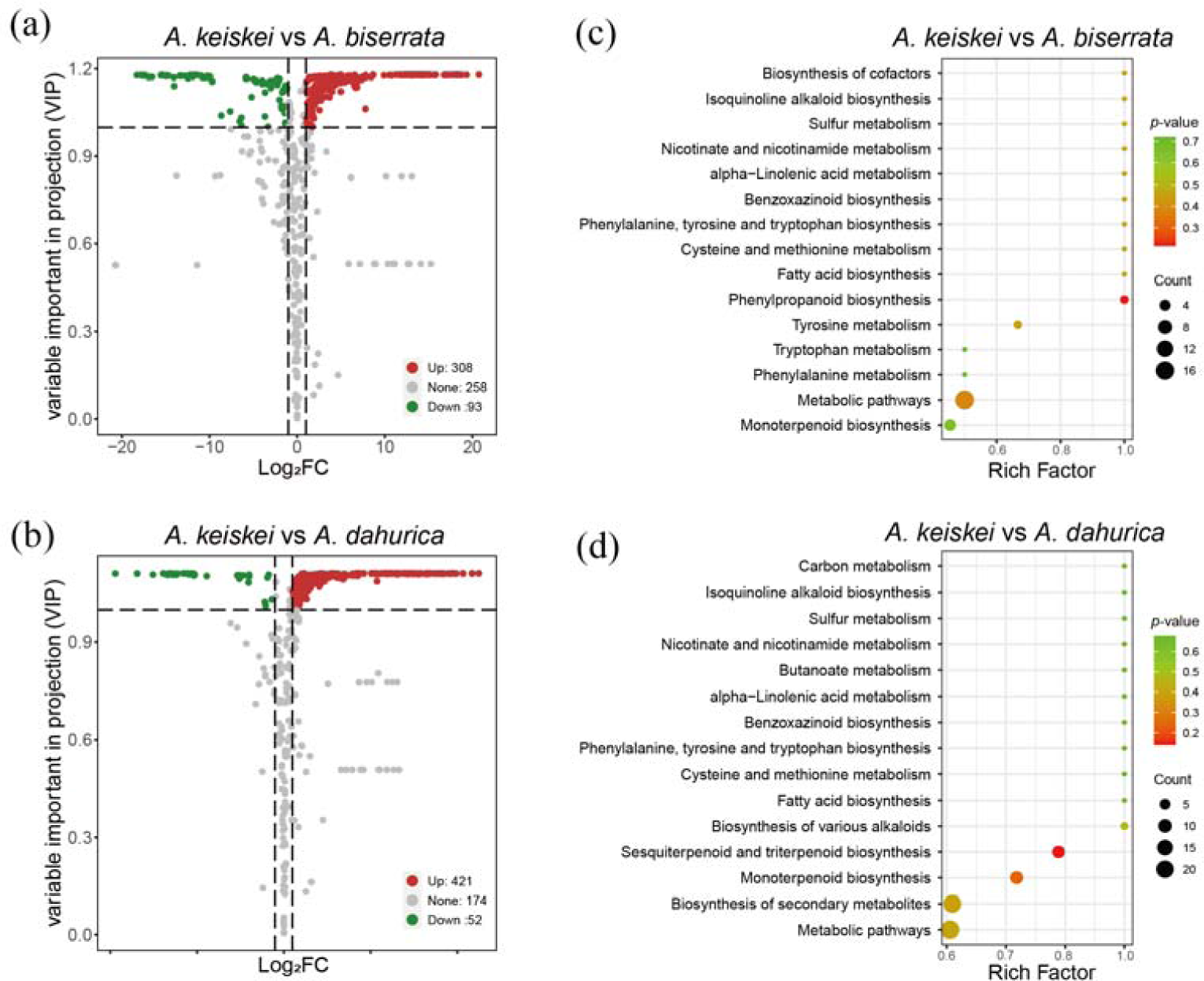
The overall distribution and KEGG enrichment analysis of differential metabolites between *A. keiskei* and *A. biserrata* (a, c), *A. keiskei* and *A. dahurica* (b, d). (a-b) Volcano plots for differential metabolites. The colors of metabolites indicated significant differences (red, upregulated; green, downregulated). (c-d) KEGG pathway enrichment analysis of differential metabolites. Color of the bubbles represented statistical significance of the enriched terms, and the size of the bubbles represented number of differential metabolites.

Moreover, to further investigate the differences of volatile metabolites in *A. keiskei* and other *Angelica* species, twenty metabolites that were differentiated the most between the two species were subsampled (Figure 8). From the comparison, the terpenoids metabolites, carvenone and cedrene were more abundant in *A. keiskei* than that in *A. biserrata*; and carene, bornyl acetate and isobornyl acetate were the most enriched in *A. keiskei* compared with *A. dahurica*. Additionally, the β-pinene was more enriched in *A. dahurica* and *A. biserrata* than in *A. keiskei*.

**Figure 8.**
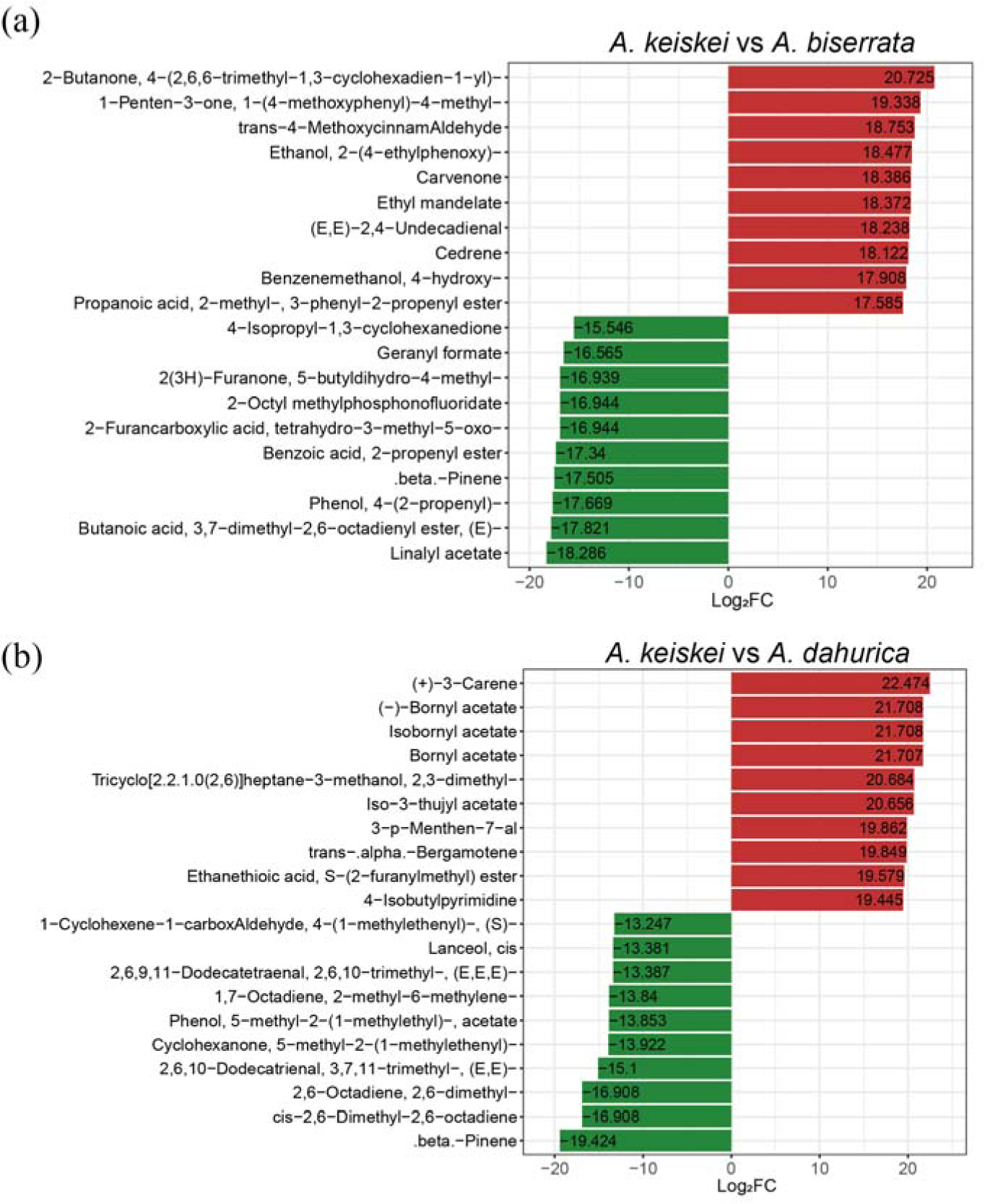
Top 20 metabolites with significant difference between *A. keiskei* and *A. biserrata* (a), *A. keiskei* and *A. dahurica* (b). Red and green represent up-regulated and down-regulated metabolites in *A. keiskei*, respectively.

## 4. Discussion

Widely targeted metabolomics offered a promising way for the chemical screening of volatile metabolites and allowed the characterization of new volatile metabolites in *Angelica* [10]. Using the method, a total of 698 volatile metabolites were identified and further classified into 15 different categories, including terpenoids, ester, heterocyclic, aromatics and 11 others (Figure 2). A clustering heat map of the metabolites showed significant difference among the four species. The number of types and abundance of volatile metabolites in *A. keiskei* was the highest. Consistent with the previous reports, terpenoids were the largest and most diverse class of volatile metabolites in the four *Angelica* species [1,2]. The pair wise comparisons between two species for the metabolite’s differences revealed that there were fewer up-regulated metabolites in *A. sinensis* when compared to the other three species (*A. dahurica, A. keiskei, A. biserrata*) whereas, relative to *A. keiskei,* most differential metabolites were down-regulated in *A. dahurica and A. biserrata.* It demonstrates that the analysis of differential metabolites is useful for understanding the differences of chemical properties among the four species.

*A. sinensis* also known as “female ginseng” is a traditional herb, which has long been used to treat various gynecological conditions [5,6,25]. Phthalides is believed to be responsible for the bioactivities of *A. sinensis* [6,26]. This study shows that 3-butylisobenzofuran-1(3H)-one and Z-ligustilide were detected in the four species and its contents were relatively higher in *A. sinensis*, which is consistent with previous studies [5]. In addition, coumarin and its derivatives are one of the important heterocyclic metabolites [27], which is mainly used as anti-HIV, anticancer activity agents, and anticoagulant activities [28,29]. The results show that the contents of 7-hydroxycoumarin and 7-ethoxycoumarin in *A. sinensis* were significantly higher than *A. dahurica*, *A. biserrata*, and *A. keiskei* (Figure 2). By virtue of its structural simplicity, 7-hydroxycoumarin has been generally accepted as the parent metabolites for the furocoumarins and pyranocoumarins and is widely used as a synthon for a wide variety of coumarin-heterocycles [30-32]. Its higher abundance in *A. sinensis* was probably associated with biosynthesis of furocoumarins and pyranocoumarins, which were reported as one of the main active components influencing the pharmaceutical activity of the herb [9]. Nevertheless, in ancient Chinese medical systems, the pharmacological effect of medicinal plants depends not only on the high abundance of a single compound, but also on the synergy of multiple active ingredients [33,34]. Furthermore, this study also found that the proportion of various components in volatile metabolites was more balanced in *A. sinensis* (Figure 2). This might explain the wide and common applications of *A. sinensis* in TCM.

Moreover, *A. keiskei* is called ashitaba in Japanese. Its leaves have been used as the medicinal part. And it is economically used as herbs, food and spices [35,36]. *A. keiskei* has been used as a medicine and food owing to its abundant pharmacological effects, including anti-cancer, lowering blood sugar and blood lipids, and improving human immunity [35,37]. However, these pharmacological effects have not been validated in scientific research. To date, it is only found in the form of raw materials in tea and cosmetics, which has limited its medicinal and clinical applications [36,38]. Interestingly, bornyl acetate, previously unmentioned terpenoid substances was detected with high expression levels in the root of *A. keiskei*, and it has been reported that bornyl acetate has antibacterial, insecticidal, and anesthetic effects symbiotically with other aromatic metabolites in the VOs [39]. This discovery provides a basis for the development and utilization of active ingredients in *A. keiskei* for health-related dietary supplements. Taken together, this study greatly enriches the database of chemical composition in *A. keiskei* and imply that *A. keiskei* exhibited benign potential to be exploited as medicinal materials and health-related dietary supplements.

Previous studies have verified that plants with closer phylogenetic relationship are not only similar in morphology but also in chemical composition and curative effects [40-42]. This study indicated the high correspondence between the volatile metabolites and the phylogenetic relationships. Although more extensive sampling and deeper investigations would be necessary to reveal more reliable correlations, the study implied that phylogenetic relationships could serve as a window to coarsely apprehend the unknown biochemical diversity of some plants based on the known biochemical map of phylogenetically related species. This finding may offer a great tool for searching replacements of medicinal plant resources that are endangered with closely related non-endangered species.

## 5. Conclusions

This study investigated the metabolites of four *Angelica* species by using widely targeted metabolomics, and found the differed accumulation of medicinally important metabolites among species. For example, high levels of bornyl acetate metabolites accumulated in *A. keiskei,* whereas coumarins and phthalides were significantly lower in *A. keiskei* than in *A. sinensis.* Moreover, the high correspondence between the dendrogram of metabolite contents and the phylogenetic tree suggested a potential correlation between the volatile metabolites and the phylogenetic relationships. Taken all together, the present study provides a biochemical map for the exploitation, application, and development of the *Angelica* species as TCM or health-related dietary supplements.

## Supplementary Materials

Figure S1: TIC chromatogram of quality control samples; Figure S2: Permutation test of OPLS-DA model; Figure S3: The violin plot of relative abundance of 15 classes in the four *Angelica* species. Table S1: The volatile metabolites detected; Table S2: The *p*-values of the t-test for 15 classes in four species.

